# Lumbar spinal Shox2 interneurons receive monosynaptic excitatory input from the lateral paragigantocellular nucleus in the adult mouse

**DOI:** 10.1101/2025.07.01.662413

**Authors:** Shayna Singh, Lihua Yao, Kimberly J. Dougherty

## Abstract

Locomotor output in vertebrates is generated in the spinal cord but is initiated and controlled by descending projections from supraspinal structures. Spinal interneurons involved in locomotion have been revealed through manipulation of genetically identified interneurons in transgenic mouse lines. Lumbar spinal interneurons expressing the transcription factor Shox2 include putative locomotor rhythm generating neurons in mice. The direct connection between supraspinal and lumbar spinal locomotor-related interneurons is comprised of reticulospinal neurons which are thought to directly provide drive to spinal rhythm generating interneurons that receive descending input and convert it to a rhythmic output. Excitatory neurons in the lateral paragigantocellular nucleus (LPGi) within the medulla have been shown to provide this descending drive in the context of forward locomotor initiation. However, a direct connection between excitatory LPGi neurons and identified spinal rhythm generating neurons has yet to be demonstrated. Here, we performed viral tracing and electrophysiological recordings to test for direct connections between the LPGi and lumbar Shox2 interneurons in adult mice. Using both monosynaptic-restricted transsynaptic rabies and anterograde AAV tracing, we show that excitatory neurons from the LPGi make direct putative excitatory synaptic contacts onto lumbar spinal Shox2 interneurons. A monosynaptic connection was confirmed via recordings of excitatory postsynaptic potentials in Shox2 interneurons in lumbar spinal slices evoked by optogenetic activation of LPGi terminals. These results demonstrate that at least a subset of lumbar spinal Shox2 interneurons receive monosynaptic excitatory input from the LPGi in the medulla, a connection which may provide the substrate for the initiation of locomotion.

## Introduction

To dynamically interact with their environments, organisms must locomote. Locomotion in vertebrates is initiated by neural circuitry spanning the entire central nervous system. A number of genetically identified populations of spinal interneurons have been shown to contribute to various aspects of locomotion (Brownstone & Bui, 2010; Dougherty, 2023). Among these, Shox2 interneurons are a putative rhythm-generating population of excitatory neurons located ventromedially in the spinal cord (Dougherty et al., 2013) exhibiting many of the criteria for rhythmogenic locomotor-related neurons (Brownstone & Wilson, 2008). Although there are other populations proposed to contribute to rhythm generation, including the Hb9 interneurons (Brownstone & Wilson, 2008; Caldeira et al., 2017; Hinckley & Ziskind-Conhaim, 2006), Lhx9 interneurons (Bertho et al., 2024), and VSCT neurons (Chalif et al., 2022), the location, electrophysiological properties, and local connectivity of the Shox2 interneurons is most consistent with a primary role in rhythm generation (Dougherty et al., 2013; Ha & Dougherty, 2018; Singh et al., 2025). However, the supraspinal structures targeting these neurons are yet unknown.

Rhythm generating neurons are hypothesized to receive an initiation signal from supraspinal structures which is converted to a rhythmic motor output (Brownstone & Wilson, 2008; Hägglund et al., 2010; Noga et al., 1988, 2003; Opris et al., 2019; Lemieux & Bretzner, 2019). The reticulospinal tract has long been suggested to be the direct link between supraspinal effectors and spinal interneurons in the context of motor output (Lloyd, 1941). The descending reticulospinal drive originating in the medulla acts as a necessary intermediary between the mesencephalic locomotor region and the spinal locomotor central pattern generator (Ausborn et al., 2019; Kim et al., 2017; Noga et al., 2003). In freely behaving mice, targeted activation within medullary reticulospinal nuclei initiates (Capelli et al., 2017) or halts (Bouvier et al., 2015; Capelli et al., 2017) locomotion. Similarly, glutamatergic reticulospinal transmission generates rhythmic locomotor-like behavior *in vitro* (Hägglund et al., 2010). Reticulospinal projections have been shown to directly contact commissural interneurons (Bannatyne et al., 2003; Matsuyama et al., 2004; Szokol et al., 2011) and motor neurons in lampreys (Ohta & Grillner, 1989). However, a determination of relation to function is complicated by the diversity in both the medullary nuclei that form the reticular spinal pathway and spinal interneuron populations.

Higher specificity has been gained more recently. Using viral tracing strategies either anterogradely to or retrogradely from specified interneuronal populations, the gigantocelluar nucleus has been shown to project to a variety of spinal neurons, including lumbar V2a interneurons (Kathe et al., 2022) involved in left-right coordination (Crone et al., 2008, 2009) and recovery of function following SCI (Kathe et al., 2022), lumbar Dmrt3 commissural neurons participating in left-right alternation (Vieillard et al., 2023), and cervical V1 interneurons (Chapman et al., 2025) mediating flexor/extensor coordination and locomotor speed (Britz et al., 2015; Gosgnach et al., 2006a). It is notable that direct LPGi input to these or other populations of locomotor-related lumbar spinal neurons have not been functionally assessed.

The lateral paragiganticellular nucleus (LPGi) has been implicated in the initiation of forward locomotion from rest (Capelli et al., 2017). Specifically, activation of glutamatergic LPGi neurons drives locomotion *in vivo* (Capelli et al., 2017). The speed of this excitatory LPGi-driven locomotor behavior was shown to scale with activation intensity (Capelli et al., 2017), suggesting that excitatory LPGi neurons have robust access to the spinal circuitry which dictates locomotor rhythm. Further, reticulospinal terminations from LPGi and neighboring caudal ventrolateral reticular nucleus forming a ‘hot spot’ for the initiation of locomotor-like activity in the spinal cord are highly dense in medial lamina VII (Hsu et al., 2023; Petras, 1967), consistent in location to be in overlap with lumbar Shox2 interneurons.

The reticulospinal tract has been the subject of studies aiming to restore motor function after spinal cord injury (Asboth et al., 2018; Baker & Perez, 2017; Kathe et al., 2022; May et al., 2017) so the identification of the specific connections made by the reticulospinal tract with spinal interneurons can be targeted in efforts to restore locomotor function after injury. Thus, our goal was to test the hypothesis that excitatory LPGi neurons make functional synaptic connections with lumbar Shox2 interneurons. We demonstrate a direct connection between excitatory neurons originating in the LPGi and lumbar spinal Shox2 interneurons in adult mice both anatomically and electrophysiologically. We show that a subset of lumbar spinal Shox2 interneurons receives excitatory monosynaptic input from the LPGi, providing evidence for a potential pathway for the initiation of locomotion.

## Materials and Methods

### Mouse Lines

All animal experiments were performed using wildtype and the following transgenic mouse lines: Shox2::Cre (Dougherty et al., 2013) and R26-lsl-tdTomato (Ai9 from The Jackson Laboratory, #007909; Madisen et al., 2010). Adult (≥P25) mice of both sex were used for this study. All experimental procedures followed National Institutes of Health guidelines and were approved by the Institutional Animal Care and Use Committee at Drexel University (LA-23-731). Mice had *ad libitum* access to food and water.

### Viral Reagents

| Virus | Addgene # | Titer | Producer |
| --- | --- | --- | --- |
| AAV9-CaMKIIa-hChR2(H134R)-EYFP | 26969 | $\geq 1 \times 10^{13}$ vg/mL | Addgene; gift from Karl Deisseroth |
| AAV9-CamKII0.4-eGFP-WPRE-rBG | 105541 | $\geq 1 \times 10^{13}$ vg/mL | Addgene; gift from James M. Wilson |
| AAV1-CAG-FLEX-H2B-HA.N2cG | 73477 | $9.35 \times 10^{12}$ GC/mL | NeuroTools Viral Vector Core, UNC; plasmid was a gift from Thomas Jessell |
| AAV1-Ef1a-FLEX-TVA-mCherry | 38044 | $1.13 \times 10^{13}$ GC/mL | NeuroTools Viral Vector Core, UNC; plasmid was a gift from Thomas Jessell |
| RabV CVS-N2c(ΔG)-eGFP | 73461 | $\geq 1.5 \times 10^8$ ffu/mL | Center for Neuroanatomy with Neurotropic Viruses (P40 OD010996) |

### Surgical Procedures

For spinal microinjections, male and female mice (P25-29) were anesthetized with isoflurane (4% induction, 2% maintenance). Dorsal skin was shaved and sterilized with betadine and isopropyl alcohol. After making an incision over lumbar spinal segments, a 1-1.5 segment laminectomy was performed, exposing the dorsal surface of the spinal cord. Four microinjections (500nL each, two on either side of the midline) of AAV1-CAG-FLEX-H2B-HA.N2cG and AAV1-Ef1a-FLEX-TVA-mCherry were delivered about 0.75mm deep into the spinal cord using a microinjection pump controller (WPI UMC4) and nanoinject II injector (Drummond 3-000-204). Following injections, dorsal skin was sutured. Mice received SR buprenorphine analgesic (0.5 mg/kg) and Baytril antibiotic (10 mg/kg) subcutaneously perisurgically. This procedure was repeated 4 weeks later in the same mice but with RabV CVS-N2c(ΔG)-eGFP (Reardon et al., 2016). CVS-N2c(ΔG) rabies microinjections (250nL each, two on either side of the midline) were delivered with the titer of ≥1.59×10^8^ ffu/mL. Mice were perfused 10 days after CVS-N2c(ΔG) rabies injection for anatomy.

For stereotaxic brainstem injections, male and female mice (>P40) were anesthetized with isoflurane (4% induction, 2% maintenance). Scalps were shaved and sterilized with betadine and isopropyl alcohol. Once secured in the stereotaxic frame (Kopf Instruments 900LS), bilateral injections (100nL each) of AAV9-CaMKIIa-hChR2(H134R)-EYFP or AAV9-CamKII0.4-eGFP-WPRE-rBG were delivered using a microinjection pump (WPI NC1987991) into the LPGi (AP - 6.96mm, ML ±0.08mm, and DV −5.7 to −6mm). Following injections, scalps were sutured. Mice received SR buprenorphine analgesic (0.5 mg/kg) and Baytril antibiotic (10 mg/kg) subcutaneously perisurgically. Mice were perfused 3 weeks following injections for anatomy or slices were prepared for electrophysiological recordings 6 weeks following injections.

### Immunohistochemistry and RNAscope in situ hybridization

Mice were anesthetized with ketamine (150mg/kg) and xylazine (15mg/kg) and perfused transcardially with 0.1M PBS, followed by 4% PFA in PBS. Spinal cords and brainstems were harvested from each animal and fixed overnight in 4% PFA solution at 4°C. Fixed tissue samples were subsequently maintained in 30% sucrose in PBS for at least 48 hours. Tissue was then embedded in OCT compound (Thermo Fisher Scientific) over dry ice and stored at −80°C. Brainstems and lumbar spinal cords were sectioned (20-40µm) transversely on a cryostat (Microm HM 505 E), directly mounted onto charged slides, and stored at −20°C. Slides were washed in PBS before being used for immunohistochemistry or RNAscope.

For immunohistochemistry, slides were first blocked in a PBS solution containing 5% donkey or goat serum, 1% bovine serum albumin, 0.2% Triton X-100, and 0.1% fish gelatin. Slides were incubated overnight in rat anti-mCherry (1:1000, Invitrogen M11217), goat anti-ChAT (1:100, Sigma AB144P), or guinea pig anti-VGLUT2 (1:200, Sigma AB2251). Slides were then incubated for 2 hours in goat anti-rat rhodamine (1:400, Invitrogen 31680), donkey anti-goat 647 secondary antibody (1:400, Jackson ImmunoResearch 705-605-003), or goat anti-guinea pig 647 secondary antibody (1:400, Invitrogen A-21450). All immunohistochemistry steps were performed at room temperature.

RNAscope was performed according to manufacturer’s protocols (Wang et al., 2012). ACDBio probes used include tdTomato (317041), eGFP (400281-C2), and Slc17a6 (456751-C3). All slides from both immunohistochemistry and RNAscope were coverslipped using Fluoromount-G with DAPI (Invitrogen 00-4959-52). Images were acquired as sequential z stacks of 20x tile-scans on a Leica DM6 fluorescence or Leica SP8 confocal microscope. eGFP RNA^+^ cell counting was performed in ImageJ using the Nucleus Counter Plugin (Collins, 2007), and overlap with VGLUT2 RNA^+^ neurons was counted manually using the multipoint tool. Puncta counting was performed manually in ImageJ using the multipoint tool. Cell area measurements were taken manually with freehand selections after adjusting image scale in ImageJ.

### Anatomical Mapping

RabV-eGFP^+^ cell bodies in medulla images were counted and marked using the multipoint tool in ImageJ and maps were constructed in Adobe Illustrator. RabV-eGFP^+^, mCherry^+^, and RabV-eGFP^+^/mCherry^+^ cell bodies in lumbar spinal cord images were counted and marked using the multipoint tool in ImageJ. Marked cell bodies were converted to representative dot maps in Adobe Illustrator. Representative dot maps were used to generate contour isoline figures in MATLAB using a custom script which is available online (https://github.com/heyshayna/SinghBrainstemManuscript2025).

### Electrophysiological Recordings

To access lumbar spinal Shox2 interneurons in the spinal slice, mice were first anesthetized with ketamine (150mg/kg) and xylazine (15mg/kg). Following decapitation and evisceration, spinal cords were removed from all mice in ice-cold dissecting solution. The dissection solution contained (in mM): 222 glycerol, 3 KCl, 11 glucose, 25 NaHCO_3_, 1.3 MgSO_4_, 1.1 KH_2_PO_4_, and 2.5 CaCl_2_. The lumbar spinal cord was sectioned transversely (300µm) in dissection solution using a vibrating microtome (Leica Microsystems). Slices were immediately transferred to recording artificial cerebrospinal fluid (ACSF) containing the following (in mM): 111 NaCl, 3 KCl, 11 glucose, 25 NaHCO_3_, 1.3 MgSO_4_, 1.1 KH_2_PO_4_, and 2.5 CaCl_2_. Slices were incubated at 34-37°C for 30 minutes and then rested at room temperature for 1 hour before recording. Dissecting and recording solutions were continuously aerated with 95%/5% O_2_/CO_2_.

Fluorescently labeled tdTomato^+^ Shox2 interneurons were visualized with a 63X objective lens on a BX51WI scope (Olympus) using LED illumination (Lumen Dynamics X-Cite) and targeted for whole cell patch clamp recordings. Electrodes were pulled to tip resistances of 5–12 MΩ using a multi-stage puller (Sutter Instruments) and were filled with intracellular solution which contained (in mM): 128 K-gluconate, 10 HEPES, 0.0001 CaCl_2_, 1 glucose, 4 NaCl, 5 ATP, and 0.3 GTP. All recordings were performed at room temperature. Data were collected with a Multiclamp 700B amplifier (Molecular Devices) and Clampex software (pClamp9, Molecular Devices). Signals were digitized at 20kHz and filtered at 6kHz. Measurements were manually calculated from recording traces in Clampfit 11 (Clampex, Molecular Devices).

Resting membrane potential was recorded shortly after gaining whole-cell access, and neurons with resting membrane potentials more depolarized than −40mV were excluded. To activate channelrhodopsin and record light-evoked postsynaptic currents, neurons were held at - 50mV in voltage clamp mode. We delivered 6 10s sweeps of 1-20Hz, 2ms pulses of 0.75-1.5mW blue LED light through the 63X objective (Lumen Dynamics X-Cite). Latency of excitatory postsynaptic currents was calculated by measuring the time between the beginning of the light stimulation artifact to the beginning of the inward current. Amplitude was measured as the maximum current response. Tetrodotoxin (0.5μM, Hello Bio HB1034) and 4-aminopyridine (100μM, Sigma 275875) were used to isolate monosynaptic light-evoked responses. Both were dissolved in recording ACSF. Values reported are averages of the first light-evoked response in each sweep.

## Results

### Transsynaptic viral-mediated tracing demonstrates a monosynaptic connection between the LPGi and Shox2 interneurons

It has been previously propounded that locomotor-related rhythmogenic spinal interneurons should receive supraspinal drive via direct excitatory reticulospinal input (Brownstone & Wilson, 2008; Hägglund et al., 2010; Noga et al., 1988, 2003; Opris et al., 2019; Lemieux & Bretzner, 2019). To determine whether a monosynaptic connection was present between the LPGi and lumbar spinal Shox2 interneurons, we performed monosynaptic-restricted transsynaptic tracing with CVS-N2c(ΔG) rabies virus (RabV CVS-N2c(ΔG)-eGFP). This strategy has previously been employed to study V1 interneurons in the cervical spinal cord (Chapman et al., 2025). We simultaneously injected two cre-dependent AAVs into the lumbar spinal cord of adult Shox2Cre mice to generate the expression of the necessary glycoprotein (N2cG) for rabies transsynaptic transmission, and the necessary cellular receptor TVA for EnvA-dependent rabies virus-host membrane fusion. We then injected CVS-N2c(ΔG) rabies virus into the same site in the lumbar spinal cord four weeks later. We found RabV-eGFP^+^/AAV1-TVA-mCherry^+^ starter cells (Fig. 1A,B) and cells which were only AAV1-TVA-mCherry^+^ (Fig. 1C) which were restricted to the ventromedial lumbar spinal cord where Shox2 interneurons reside. However, cells which were only RabV-eGFP^+^ extended throughout the dorsal horn in the lumbar spinal cord as well (Fig. 1D).

**Figure 1:**
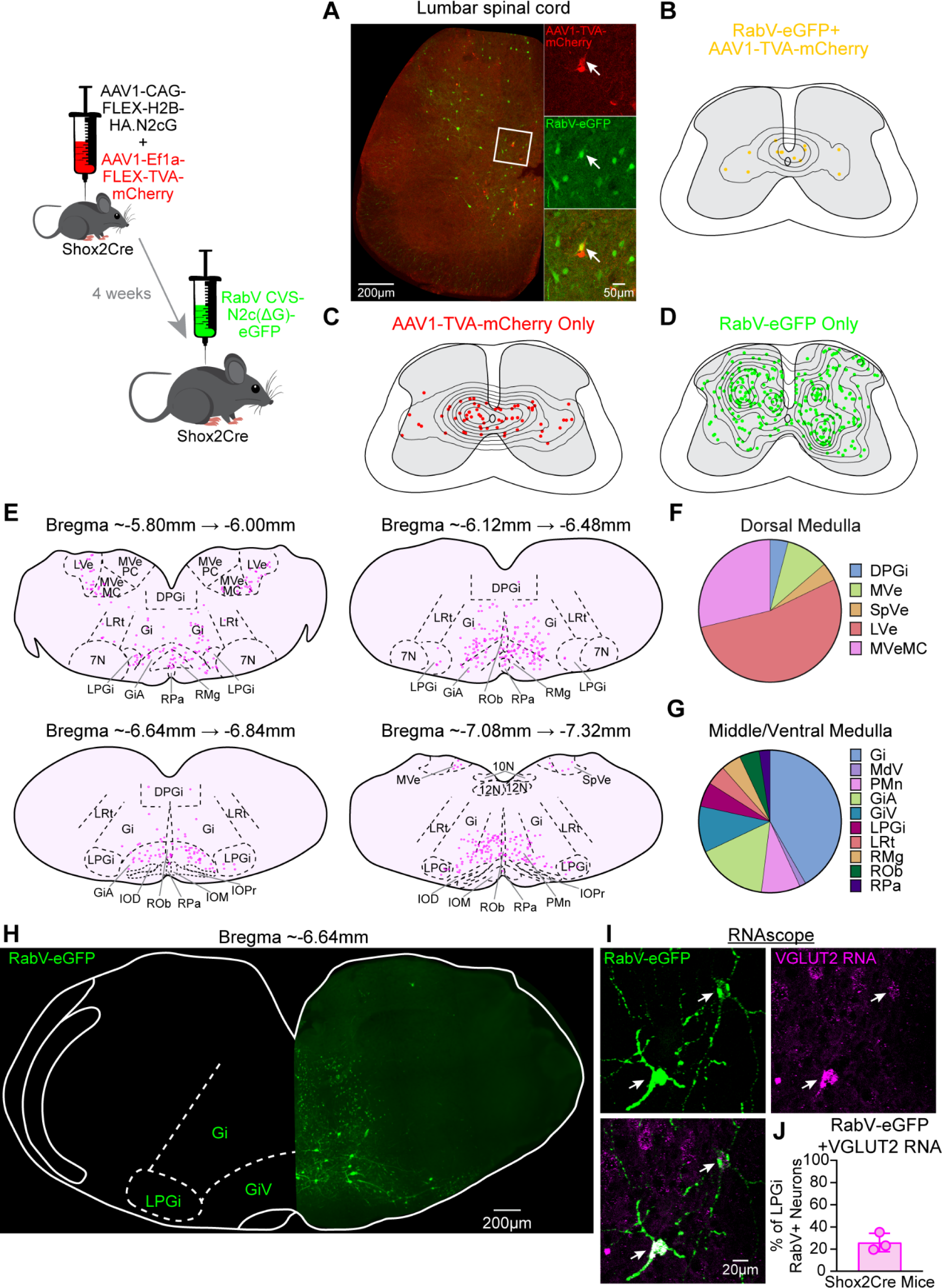
Transsynaptic rabies tracing from lumbar spinal Shox2 interneurons reveals starter cells and monosynaptically-connected cells in the lumbar spinal cord. (A) Representative image following bilateral microinjections of helper AAVs, AAV1-CAG-FLEX-H2B-HA.N2cG and AAV1-Ef1a-FLEX-TVA-mCherry, and RabV CVS-N2c(ΔG)-eGFP showing RabV-eGFP^+^ cells (green), AAV1-TVA-mCherry^+^ cells (red), and RabV-eGFP^+^/AAV1-TVA-mCherry^+^ starter cells (yellow) present in the lumbar spinal cord. (B) Density contour plot of mapped RabV-eGFP^+^/AAV1-TVA-mCherry^+^ starter cell bodies. (C) Density contour plot of mapped AAV1-TVA-mCherry^+^ cell bodies. (D) Density contour plot of mapped RabV-eGFP^+^ cell bodies. (E) eGFP^+^ cells are present in the medulla (green) at approximately Bregma −6.64. (F,G) Cell counting in the LPGi after probing for eGFP RNA and VGLUT2 RNA using RNAscope demonstrates that approximately 22% of eGFP RNA^+^ cells (green) are also VGLUT2 RNA^+^ (magenta). (H) Representative maps of cell body positions of RabV-eGFP^+^ cells. (I) Relative distribution of RabV-eGFP^+^ cells within nuclei in the dorsal medulla. Data represent the proportion of the total eGFP+ cell count in the dorsal medulla. (J) Relative distribution of RabV-eGFP+ cells within nuclei in the middle and ventral medulla. N=3 mice. DPGI, dorsal paragigantocellular nucleus; MVe, medial vestibular nucleus; SpVe, spinal vestibular nucleus; LVe, lateral vestibular nucleus; MVeMC, medial vestibular nucleus magnocellular part; Gi, gigantocellular nucleus; MdV, ventral medial reticular nucleus; PMn, paramedian reticular nucleus; GiA, anterior gigantocellular nucleus; GiV, ventral gigantocellular nucleus; LPGi, lateral paragigantocellular nucleus; LRt, lateral reticular nucleus; RMg, raphe magnus; ROb, raphe obscuruis; RPa, raphe pallidus (Paxinos & Franklin, 2004).

We mapped RabV-eGFP^+^ neurons in sections containing the reticulospinal nuclei from the medulla (Fig. 1E). We found eGFP^+^ neurons in the LPGi, gigantocellular nucleus, ventral gigantocellular nucleus, anterior gigantocellular nucleus, the medullary reticular formation ventral part, the paramedian reticular nucleus, the lateral reticular nucleus, and several raphe nuclei (Fig. 1E,F). Dorsal medullary nuclei also contained RabV-eGFP^+^ cells, including the dorsal paragigantocellular nucleus and various vestibular nuclei (Fig. 1E,G).

We then focused on the LPGi neurons which monosynaptically contact lumbar spinal Shox2 interneurons (Fig. 1H). We performed RNAscope (Fig. 1I) which revealed that a subset (22%) of eGFP^+^ neurons in the LPGi which monosynaptically contact lumbar spinal Shox2 interneurons contain VGLUT2 RNA (Fig. 1J). This demonstrates that excitatory neurons in the LPGi are monosynaptically connected to lumbar spinal Shox2 interneurons.

We performed control experiments in age-matched wildtype mice which do not express cre. Injections of only RabV-eGFP (Supplementary Fig. 1A) or AAV1-CAG-FLEX-H2B-HA.N2cG, AAV1-Ef1a-FLEX-TVA-mCherry, and RabV-eGFP (Supplementary Fig. 1B) resulted in no fluorescently labeled cells in either the spinal cord or in the medullary nuclei. Any fluorescence detected in these images was auto-fluorescence and was equally intense in all testable channels. Cell counts from experimental and control injections in all examined regions are compiled in Supplementary Table 1. Taken together, these data demonstrate that lumbar spinal Shox2 interneurons receive monosynaptic connections from many medullary nuclei and spinal cord cells, including excitatory LPGi neurons.

### Anterograde viral-mediated tracing demonstrates a connection between excitatory LPGi neurons and Shox2 interneurons

To validate the connection between the LPGi and lumbar spinal Shox2 interneurons, anterograde viral-mediated tracings were performed. This strategy efficiently labels the LPGi and reveals the connections to all spinal neurons, not specifically to the Shox2 interneurons. Following bilateral injections of AAV9-CamKII0.4-eGFP into the LPGi in adult Shox2Cre;tdTomato mice (Fig. 2A), we examined projections and terminations in lumbar spinal cord sections (Fig. 2B). The distribution of eGFP in the spinal cord matches previous studies examining activation patterns following LPGi stimulation (Hsu et al., 2023), with hot spots of descending fibers within the lateral white columns and dense terminations medially in spinal gray matter. These findings also match what has been shown in anatomical tracing studies of the reticulospinal tract, originating from the LPGi, in adult mice (Liang et al., 2016).

**Figure 2:**
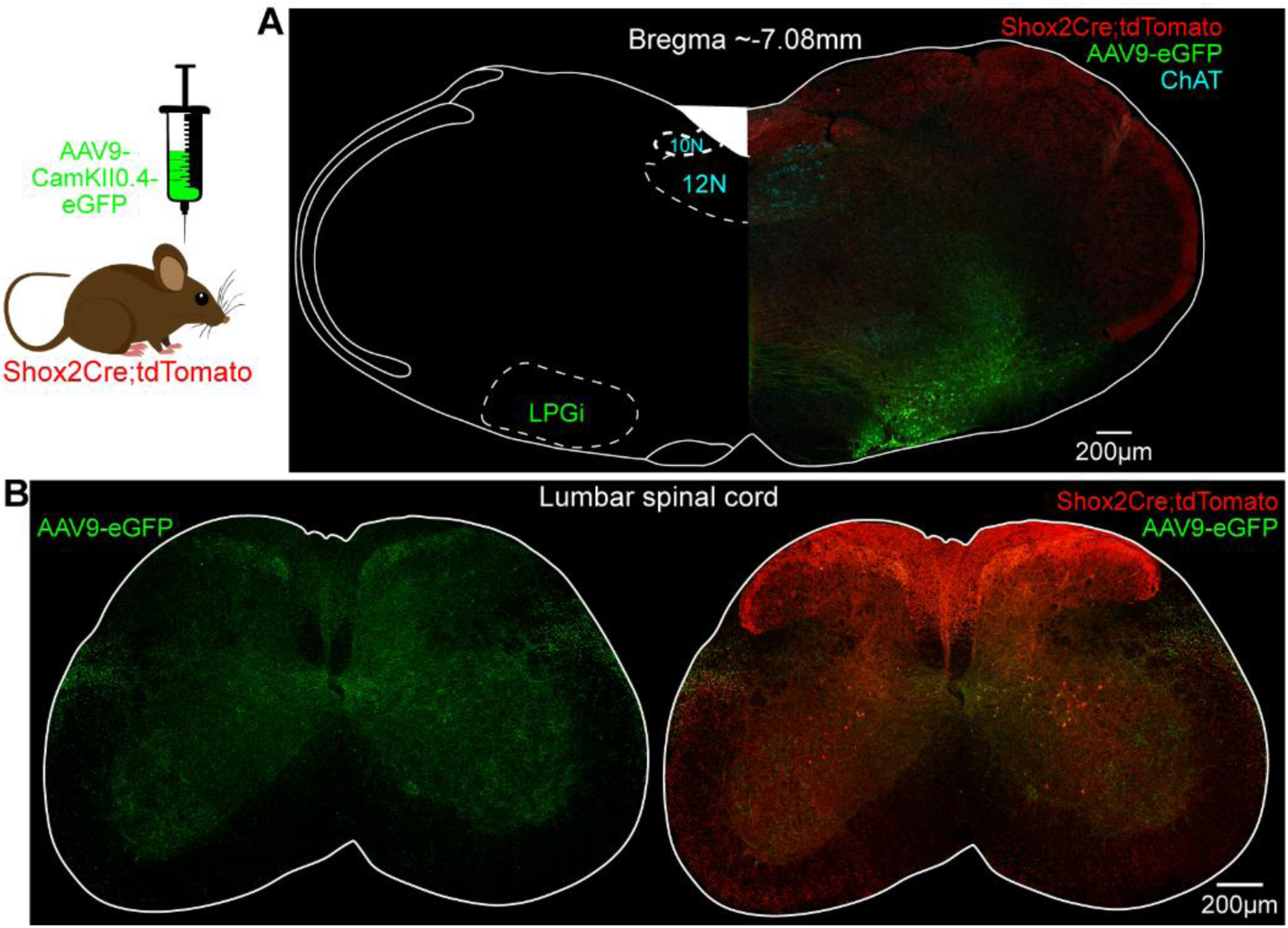
Anterograde viral tracing of LPGi spinal projections. (A) Bilateral injections of AAV9-CamKII0.4-eGFP-WPRE-rBG were delivered to adult Shox2Cre;tdTomato mice, resulting in fluorescently labeled cell bodies in the LPGi (green) at the level of the ChAT^+^ 10^th^ and 12^th^ cranial nerve motor nuclei (cyan) at approximately −7.08 from Bregma (Paxinos & Franklin, 2004). (B) eGFP+ projections in the lumbar spinal cord (green) overlap with the ventromedial location of Shox2 interneurons (red). N=4 mice.

The LPGi is constituted of serotonergic, glutamatergic, GABAergic, and glycinergic neurons (Capelli et al., 2017; Ganley et al., 2023; Hsu et al., 2023; Talluri et al., 2022). However, the activation of excitatory LPGi neurons, and not glycinergic or GABAergic neurons, was shown to promote forward locomotion in adult mice (Capelli et al., 2017). We therefore sought to calculate the proportion of transfected LPGi neurons which are excitatory. We probed for eGFP RNA to visualize transfected neurons, and VGLUT2 RNA to visualize excitatory neurons (Fig. 3A). We found that an average of 42% eGFP RNA^+^ LPGi neurons were VGLUT2 RNA^+^ across four mice (Fig. 3B,C). This suggested that a proportion of excitatory neurons were targeted by our injections into the LPGi. To examine the terminations of these neurons onto lumbar spinal Shox2 interneurons, we performed immunohistochemistry on spinal cord slices to label VGLUT2^+^/eGFP^+^ puncta in apposition to Shox2 interneurons (Fig. 3D). We show that, while a subset of lumbar Shox2 interneurons had no VGLUT2^+^/eGFP^+^ puncta (Fig. 3E), most Shox2 interneurons were 50-200μm^2^ in area and overlapped with 10 or less VGLUT2^+^/eGFP^+^ puncta (Fig. 3F). Some Shox2 interneurons were considerably larger (300-750 μm^2^) and contained higher amounts of overlapping VGLUT2^+^/eGFP^+^ puncta (Fig. 3F), supporting the notion that lumbar spinal Shox2 interneurons are a heterogeneous population. Moreover, almost half of the eGFP^+^ puncta on lumbar spinal Shox2 interneurons were VGLUT2^+^ (Fig. 3G). This anatomical data supports that Shox2 interneurons receive input from descending LPGi neurons, with at least half of it being from excitatory neurons in the LPGi.

**Figure 3:**
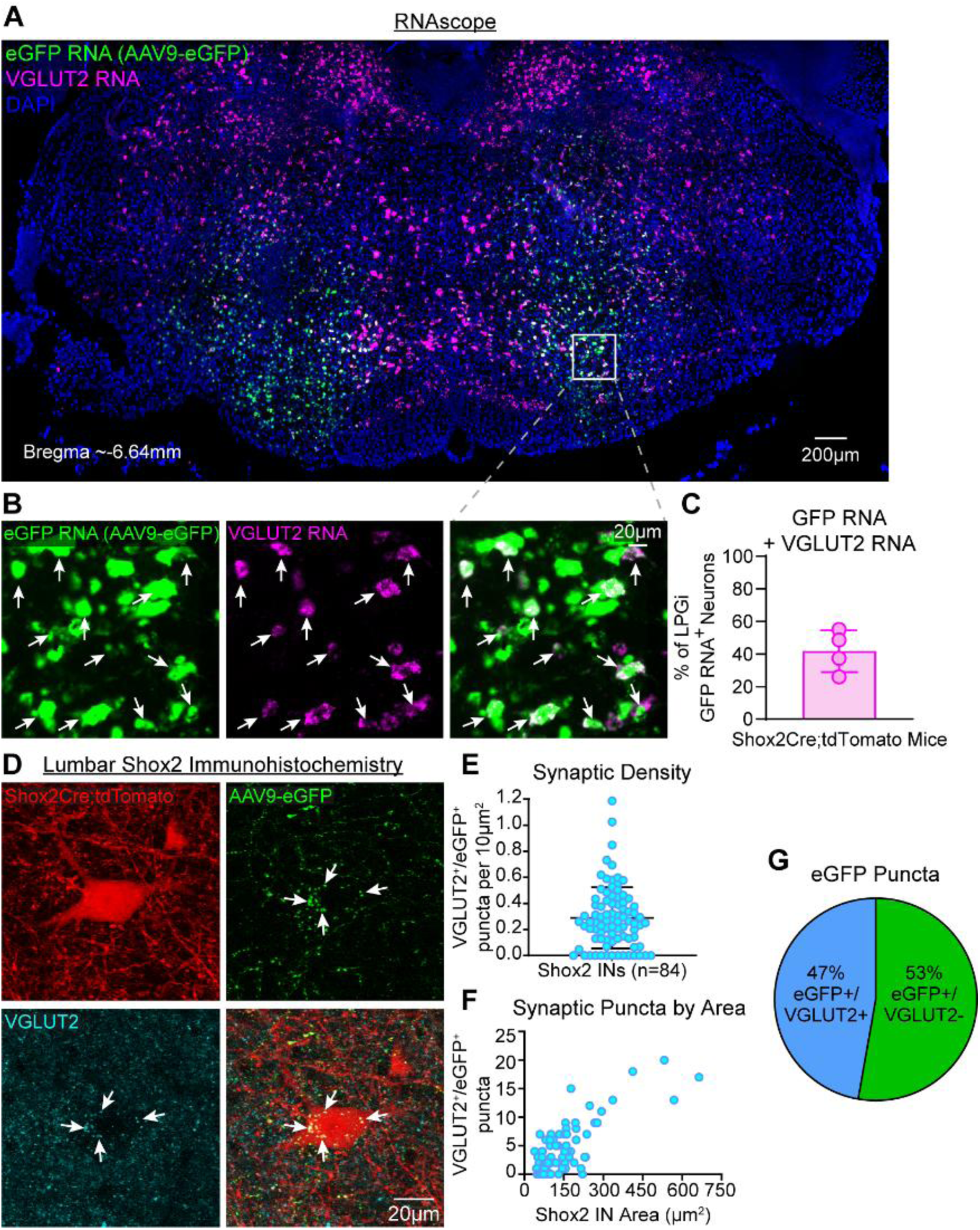
RNAscope in LPGi and immunohistochemistry in lumbar spinal cord reveal excitatory anatomy. (A) RNAscope showing bilateral AAV9-CamKII0.4-eGFP injection (green) and glutamatergic cells signified by VGLUT2 RNA (magenta) with DAPI for whole-slice morphology of brainstem slice (blue) at approximately Bregma −6.64 (Paxinos & Franklin, 2004). (B) Inset of eGFP RNA (green), VGLUT2 RNA (magenta) and overlapping image showing putatively excitatory LPGi neurons. (C) Approximately 40% of injected, eGFP RNA^+^, LPGi neurons also are VGLUT2 RNA^+^. (D) Staining lumbar spinal slices from AAV9-eGFP injected mice for VGLUT2 (cyan) shown overlap with eGFP^+^ terminations (green) onto Shox2 interneurons (magenta). (E) The density of putatively glutamatergic synaptic contacts from the LPGi onto lumbar spinal Shox2 interneurons. (F) Number of putatively glutamatergic synaptic contacts plotted against Shox2 interneuron cross-sectional area. (G) Of all eGFP+ puncta counted on Shox2 interneurons, 47% were also VGLUT2^+^. n=84 lumbar shox2 interneurons, N=4 mice.

### Electrophysiology demonstrates an excitatory monosynaptic connection between the LPGi and lumbar spinal Shox2 interneurons

In order to evaluate the functionality of the connection between the LPGi and Shox2 interneurons in the adult mouse, we sought to directly measure the electrophysiological input that Shox2 interneurons receive via these synaptic contacts. To determine the extent to which the observed putative excitatory puncta were functional synapses, we performed bilateral injections of AAV9-CaMKIIa-hChR2(H134R)-EYFP into the LPGi in adult Shox2Cre;tdTomato mice (Fig. 4A). We optically stimulated LPGi terminals in the lumbar spinal slice during whole-cell patch clamp from visually-identified Shox2 interneurons. We did this in both baseline recordings and in the presence of TTX+4-AP to isolate the monosynaptic component of the light-evoked response (Fig. 4B). We found that one third (n=12/36) of tested Shox2 interneurons displayed baseline light-evoked excitatory postsynaptic currents (EPSCs, Fig. 4C). Of this subset, we lost three Shox2 interneurons and were therefore unable to confirm whether the input they received was monosynaptic or polysynaptic. Following TTX+4-AP application, we determined that a subset of Shox2 interneurons received monosynaptic light-evoked input (n=4/9, Fig. 4F). There was no significant change in EPSC amplitude (16.7pA ± 6.7pA) or latency (6.5ms ± 1.5ms) in Shox2 interneurons with detectable inputs remaining, compared to baseline amplitude (25.0pA ± 8.8pA; Fig. 4G) and latency (4.9ms ± 0.7ms; Fig. 4H). The mean amplitude of the confirmed-polysynaptic EPSCs recorded in 5 Shox2 interneurons at baseline (23.8pA ± 7.8pA, Fig. 4D) was similar to the mean monosynaptic EPSC amplitude at baseline (25.0pA ± 8.8pA, Fig. 4G). The mean amplitude of the unconfirmed-polysynaptic Shox2 interneurons was 37.1pA ± 3.3pA (Fig. 4D). The mean latency of the confirmed-polysynaptic input to Shox2 interneurons (6.0ms ± 1.9ms, Fig. 4E) was also similar to the mean baseline monosynaptic EPSC latency (4.9ms ± 0.7ms, Fig. 4H) and unconfirmed-polysynaptic Shox2 interneuron mean latency (5.8ms ± 1.9ms, Fig. 4E). Taken together, these findings demonstrate that a subset of lumbar spinal Shox2 interneurons receive monosynaptic excitatory input from the LPGi.

**Figure 4:**
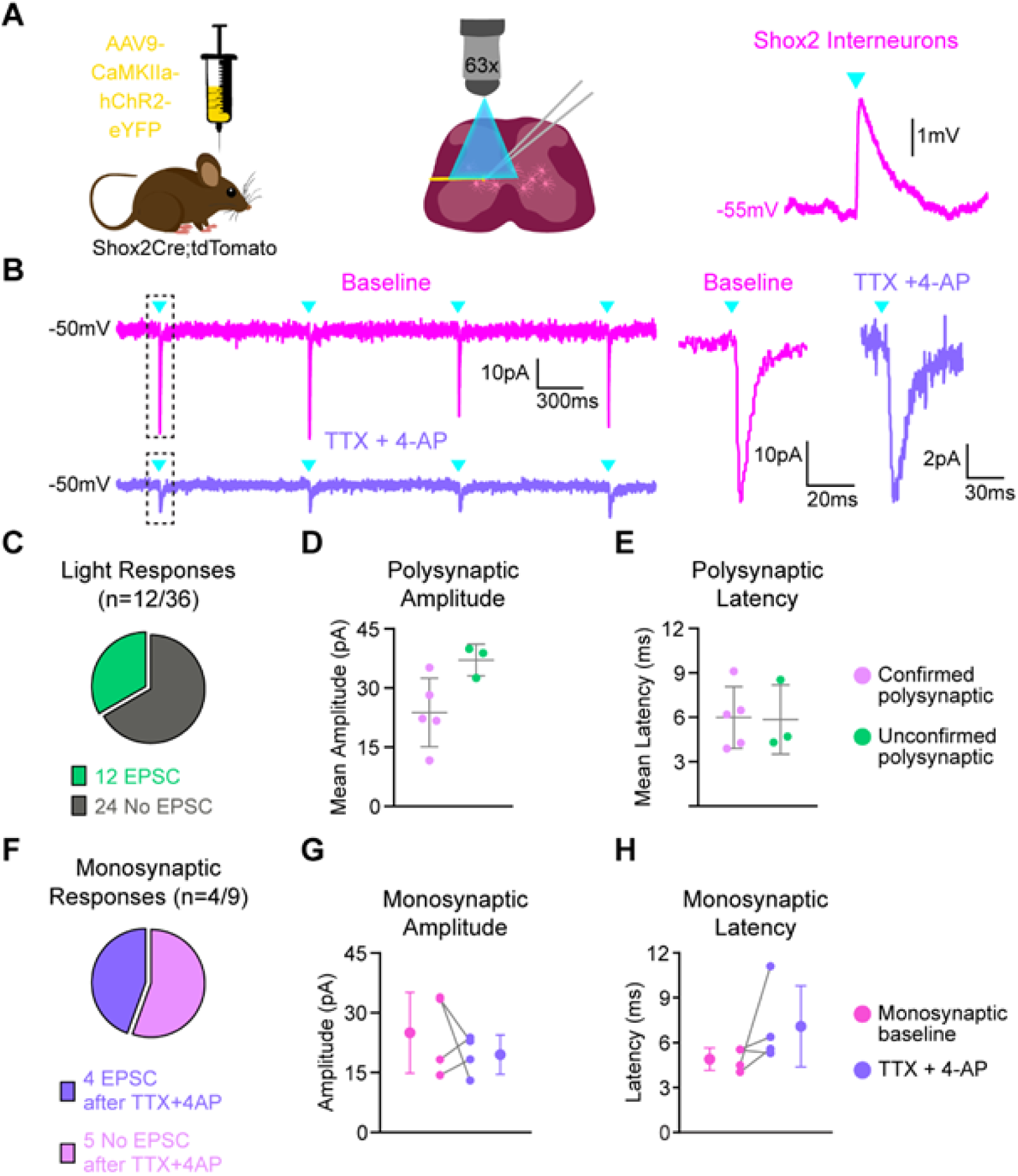
Whole-cell patch clamp recordings of light-evoked LPGi terminal activation in Shox2 interneurons. (A) Bilateral injections of AAV9-CaMKIIahChR2-eYFP were delivered to adult Shox2Cre;tdTomato mice. Through the microscope objective, fluorescent light evoked activation of ChR2 in LPGi terminals in the lumbar spinal slice during whole-cell patch clamp of Shox2 interneurons. In current-clamp mode, light evoked excitatory post-synaptic potentials. (B) Light pulse trains evoked excitatory postsynaptic currents in Shox2 interneurons in baseline ACSF (magenta), which had a monosynaptic component revealed after the bath application of TTX + 4-AP (purple). (C) Of the 36 Shox2 interneurons recorded from, 12 displayed light-evoked excitatory postsynaptic potentials in baseline ACSF conditions. 3 of the 12 interneurons with responses were lost after baseline recordings. (D) Amplitude and (C) latency of 3 unconfirmed-polysynaptic (green) and 5 confirmed-polysynaptic (pink) Shox2 interneurons. (F) Of the 9 Shox2 interneurons with light-evoked EPSCs that were tested further, 4 had light-evoked EPSCs which persisted following bath application of TTX + 4-AP. (G) Amplitude and (H) latency of 4 monosynaptic Shox2 interneurons in baseline (purple) and drug (magenta) conditions. N=7 mice.

## Discussion

Using anatomy and electrophysiology, we demonstrated that a subset of Shox2 interneurons receive monosynaptic excitatory input from the LPGi in the adult mouse. This study supports that Shox2 interneurons may be an entry point for supraspinal descending drive into spinal locomotor circuitry.

### Technical considerations of viral tracing and electrophysiological connectivity testing

We first sought to examine the monosynaptic component of the connection between the LPGi and Shox2 interneurons to corroborate predictions of locomotor-related brainstem centers sending descending drive directly to rhythm-related spinal interneurons (Ausborn et al., 2019; Kim et al., 2017; Noga et al., 2003). We chose the CVS-N2c(ΔG) rabies virus-mediated strategy, which has been previously used in mice to examine monosynaptic inputs to V1 interneurons (Chapman et al., 2025). The use of the CVS-N2c(ΔG) rabies virus has advantages over earlier strains in that it allows for the efficient and intense labeling of neuronal somas and processes over long distances in a G protein-dependent/monosynaptic-restricted fashion (Reardon et al., 2016; Wickersham, Finke, et al., 2007; Wickersham, Lyon, et al., 2007). However, it is impossible to determine the true number of starter neurons due to the toxicity of the CVS-N2c(ΔG) rabies virus (Reardon et al., 2016). Further, the starter population is likely to be an underestimation or a small subset of the local Shox2 interneuron population due to the necessity of three essential components (TVA, G, and N2c(ΔG)) to be successfully expressed in the same neuron. Nevertheless, we were able to identify labeled spinal starter neurons and synaptically coupled neurons in the spinal cord and brainstem.

Anterograde AAV tracing was used to complement the retrograde tracing and to examine the termination pattern of the LPGi reticulospinal neurons in the lumbar spinal cords of adult mice. Our findings are comparable to what has been previously described in the neonatal *in vitro* preparation (Hsu et al., 2023) and in the adult mouse using biotinylated dextran amine solution (Liang et al., 2016). We chose the CaMKIIα promoter for its specificity to supraspinal excitatory neurons (Yizhar et al., 2011). More recently however, it has been shown that viral labeling using CaMKIIα promoters is efficient in both excitatory and inhibitory supraspinal neurons (Veres et al., 2023). Our findings support this, as we demonstrate that only about half of the targeted cells in the LPGi labeled in our anterograde AAV tracing experiments are VGLUT2 RNA^+^. The LPGi includes serotonergic, glutamatergic, GABAergic, and glycinergic neurons (Capelli et al., 2017; Ganley et al., 2023; Hsu et al., 2023; Talluri et al., 2022). None of these discrete neuronal types in the LPGi have identified spinal targets beyond the excitatory connections to Shox2 interneurons presented in this study. Moreover, it is likely that LPGi inputs to different subsets of Shox2 and other spinal interneurons are serotonergic, excitatory, and inhibitory.

Only excitatory light-evoked responses from the LPGi in the spinal slice were recorded in this study. Electrophysiological testing may underestimate the connections between the targeted LPGi neurons and Shox2 interneurons for many reasons. We may be eliminating some of the synaptic connections onto Shox2 interneuron during slice collection due to the removal of dendritic arbors which extend away from the Shox2 interneuron soma beyond the range of each 300µm slice. We are also unable to determine how many synaptic contacts are necessary for the light-evoked EPSCs. The amplitude of the light evoked EPSCs is also likely dictated by light intensity, and ours was relatively low compared to similar experiments (Chapman et al., 2025). Furthermore, the expected response latency at room temperature is unknown.

### Adult lumbar Shox2 interneurons likely integrate information broadly from reticular and local spinal sources

Experiments aimed at determining the function of genetically-identified locomotor circuit interneurons have largely been carried out in neonatal animals (Crone et al., 2008; Gosgnach et al., 2006b; Lanuza et al., 2004), with some exceptions (Crone et al., 2009; Koronfel et al., 2021; Talpalar et al., 2013). This is particularly true of rhythm generating populations, since most of the manipulations affect respiratory function and/or feeding, limiting viability (Bertho et al., 2024; Caldeira et al., 2017; Dougherty et al., 2013). This is in contrast with the majority of experiments which determined the roles of various reticulospinal populations. These were largely performed in adult mice, due to the use of viral tools, which require weeks to express (Bouvier et al., 2015; Capelli et al., 2017; Cregg et al., 2020; Esposito et al., 2014; Talpalar et al., 2013; Usseglio et al., 2020). The direct excitatory connections demonstrated in this study are likely already present at birth, as reticulospinal input develops embryonically (Perreault & Glover, 2013). Further, descending fiber-evoked locomotion is reduced in frequency when Shox2 neurons are synaptically silenced (Dougherty et al., 2013) and stimulation of the LPGi induces locomotor-like activity (Hsu et al., 2023) in the reduced neonatal mouse preparation. Thus, it is possible that prior demonstrations of reticulospinal activation evoking locomotor-like activity in neonatal preparations (Hägglund et al., 2010; Hsu et al., 2023) are via this pathway.

Although the identities of the spinal neurons presynaptic to Shox2 interneurons were not explored, the locations of the presynaptic neurons offer hints to the spinal circuit architecture. Local presynaptic neurons were found in most laminae but there were concentrations in the deep dorsal horn and in the medial ventral horn (lamina VIII). Commissural interneurons are concentrated in lamina VIII (Eide et al., 1999; Jankowska & Noga, 1990). At least a subset of Shox2 interneurons activate commissural interneurons (Dougherty et al., 2013) but the connection between commissural neurons and Shox2 (or rhythm generating) interneurons is predicted to be important for the coordination of left and right sides during locomotion (Shevtsova et al., 2015). Deep dorsal presynaptic neurons would be consistent with those in reflex pathways to rhythm generating neurons (Domínguez-Rodríguez et al., 2020; Frigon et al., 2022), which have been shown to be both excitatory and inhibitory to Shox2 interneurons in both neonate (Li et al., 2019) and adult (Garcia-Ramirez et al., 2021). Other local neurons in lamina VII are also expected to be connected, including other Shox2 neurons (Ha & Dougherty, 2018) in addition to V1 and V2b inhibitory interneurons involved in flexor-extensor alternation (Britz et al., 2015; Shevtsova et al., 2022; Shevtsova & Rybak, 2016). The direct testing of the connectivity of these populations is complicated by the downregulation of the expression of the identifying transcription factors. It is clear that Shox2 interneurons broadly receive input from spinal neurons, which may be integrated into their rhythmogenic output in locomotor circuitry.

### Potential behavioral implications including the initiation of forward locomotion

In the mammalian medullary reticular formation, regional borders are ambiguous and neuronal cell types are diverse (Ascenzi, 2025). Robust behavioral effects after the manipulation of specific medullary neuronal populations have been evoked in many motor-related contexts in mice, including ipsilateral body turning (Cregg et al., 2020), locomotor arrest (Bouvier et al., 2015), REM sleep and associated muscle atonia (Weber et al., 2015), and even wakefulness with strong postural tone from coma (Gao et al., 2019). Based on the monosynaptically-connected medullary nuclei identified in this study, Shox2 interneurons may play a role in these and other motor functions. It is possible that subsets of lumbar spinal Shox2 interneurons play regulatory roles in posture and gait, as suggested by their direct input from vestibular nuclei such as the lateral vestibular nucleus (Witts & Murray, 2019). Shox2 interneurons may also directly integrate raphespinal input in the context of locomotion, as the caudal raphe nuclei including the raphe pallidus have been implicated in forward locomotion (Jordan, 1998). The nature and utility of these connections to lumbar spinal Shox2 interneurons are likely varied and relevant to broad arrays of behavioral outcomes.

The LPGi alone is implicated in muscle atonia and postural control in addition to locomotion, with the neurons controlling these behaviors likely intermingled within this region (Perreault & Giorgi, 2019). Other functions the LPGi has been implicated in include sexual reflexes in male rodents (Liu & Sachs, 1999; Marson et al., 1992), audition (Bellintani-Guardia et al., 1996), pain (Ganley et al., 2023), bladder control (Talluri et al., 2022), and cardiac function (Dergacheva et al., 2010). Previous studies have also demonstrated the role of the LPGi in general arousal (Guyenet et al., 1990; Van Bockstaele & Aston-Jones, 1995; Zhang et al., 2024). This is likely via the dense connections from the LPGi to the locus coeruleus which are comprised of excitatory and inhibitory projection neurons and have been shown to collateralize to innervate the spinal cord as well (Van Bockstaele & Aston-Jones, 1995). It is unclear how many simultaneously ascending and descending neurons exist in reticular nuclei, let alone the LPGi (Pfaff et al., 2012). Lumbar spinal Shox2 interneurons may integrate excitatory and inhibitory input from descending bifurcating neurons to modulate features of locomotor behavior.

It is clear that the effect of direct unilateral activation of excitatory LPGi neurons is the initiation of locomotion, which can be driven in a speed-dependent manner (Capelli et al., 2017), and has been reproduced using computational modeling (Ausborn et al., 2019). As putative rhythm-generating neurons of the central pattern generator and direct recipients of excitatory reticulospinal input from the LPGi, spinal Shox2 interneurons are poised to mediate these effects. The identities and locations of the excitatory LPGi neurons necessary for the initiation of locomotion, and what proportion of these are present in this study as monosynaptic partners to Shox2 interneurons is unknown. Future investigations are also needed to delineate the utility of the connection between the LPGi and lumbar spinal Shox2 interneurons in the context of motor and other behaviors, including and beyond the excitatory connections identified in this study.

## Acknowledgements

We thank members of the Marion Murray Spinal Cord Research Center and Drexel ULAR for support. We thank Rémi Ronzano, Mary Patton, Anand Kulkarni, and Jay Bikoff for experimental advice. We thank Julien Bouyer, Ying Jin, Wenqiang Huang, and Dong Wang for generous surgical assistance. We also thank Malcolm Jennings for experimental assistance. This work was supported by NIH R01 NS130799 (KJD), T32 NS121768 (SS), and F31 NS132514 (SS). Modified rabies virus was provided by the NIH Virus Center supported by P40 OD010996.

