## Supplemental information for "Lumbar spinal Shox2 interneurons receive monosynaptic excitatory input from the lateral paragigantocellular nucleus in the adult mouse"

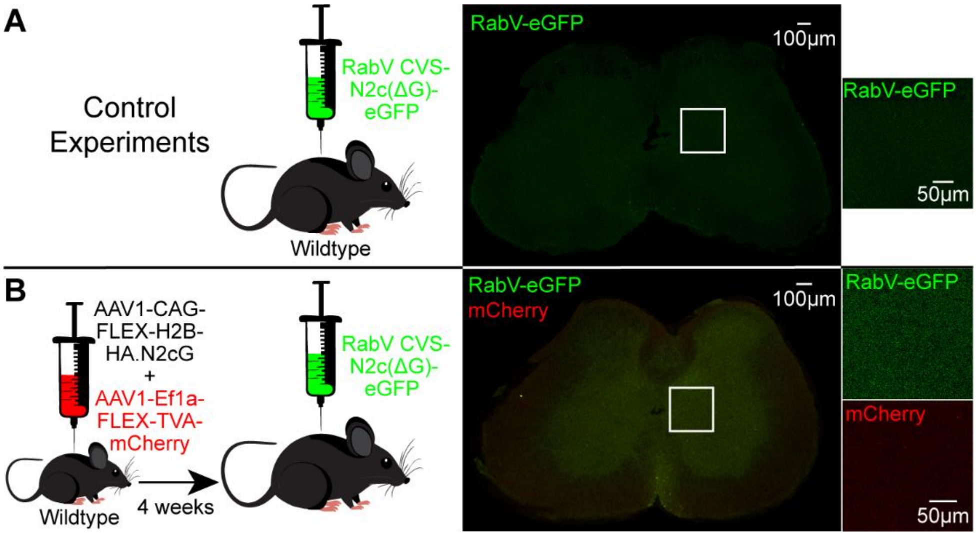


**Supplemental Figure 1: Control experiments in wildtype mice demonstrate virus specificity.** (A) Bilateral RabV CVS-N2c(ΔG)-eGFP microinjections in adult wildtype mice resulted in no fluorescently labeled cells in the lumbar spinal cord. N=2 mice. (B) Bilateral microinjections of AAV1-CAG-FLEX-H2B-HA.N2cG+AAV1-Ef1a-FLEX-TVA-mCherry and RabV CVS-N2c(ΔG)-eGFP in adult wildtype mice resulted in no fluorescently labeled cells in the lumbar spinal cord. N=3 mice.

**Supplemental Table 1: Viral tracing cell counts by region**

| Region | **Shox2Cre N=3** AAV1-N2cG+AAV1-TVA-mCherry+RabV-eGFP into lumbar spinal cord | | | | **Wildtype N=2** RabV-eGFP into lumbar spinal cord | | **Wildtype N=3** AAV1-N2cG+AAV1-TVA-mCherry+RabV-eGFP into lumbar spinal cord | | |
|---|---|---|---|---|---|---|---|---|---|
| | **Mouse 1** | **Mouse 2** | **Mouse 3** | **Mean** | **Mouse 1** | **Mouse 2** | **Mouse 1** | **Mouse 2** | **Mouse 3** |
| Lumbar Spinal Cord mCherry only | 22 | 69 | 41 | 44 | 0 | 0 | 0 | 0 | 0 |
| Lumbar Spinal Cord eGFP only | 167 | 61 | 430 | 219 | 0 | 0 | 0 | 0 | 0 |
| Lumbar Spinal Cord mCherry+eGFP | 7 | 1 | 9 | 6 | 0 | 0 | 0 | 0 | 0 |
| DPGi eGFP | 2 | 0 | 1 | 1 | 0 | 0 | 0 | 0 | 0 |
| Mve eGFP | 1 | 2 | 4 | 2 | 0 | 0 | 0 | 0 | 0 |
| SpVe eGFP | 0 | 0 | 3 | 1 | 0 | 0 | 0 | 0 | 0 |
| LVe eGFP | 19 | 2 | 0 | 7 | 0 | 0 | 0 | 0 | 0 |
| MVeMC eGFP | 8 | 0 | 13 | 7 | 0 | 0 | 0 | 0 | 0 |
| Gi eGFP | 103 | 12 | 89 | 68 | 0 | 0 | 0 | 0 | 0 |
| MdV eGFP | 0 | 0 | 7 | 2 | 0 | 0 | 0 | 0 | 0 |
| PMn eGFP | 0 | 8 | 35 | 14 | 0 | 0 | 0 | 0 | 0 |
| GiA eGFP | 50 | 0 | 28 | 26 | 0 | 0 | 0 | 0 | 0 |
| GiV eGFP | 12 | 10 | 29 | 17 | 0 | 0 | 0 | 0 | 0 |
| LRt eGFP | 0 | 2 | 8 | 3 | 0 | 0 | 0 | 0 | 0 |
| RMg eGFP | 12 | 0 | 10 | 7 | 0 | 0 | 0 | 0 | 0 |
| ROb eGFP | 7 | 1 | 14 | 7 | 0 | 0 | 0 | 0 | 0 |
| RPa eGFP | 8 | 0 | 4 | 4 | 0 | 0 | 0 | 0 | 0 |
| LPGi eGFP only | 18 | 0 | 6 | 8 | 0 | 0 | 0 | 0 | 0 |

DPGI, dorsal paragigantocellular nucleus; MVe, medial vestibular nucleus; SpVe, spinal vestibular nucleus; LVe, lateral vestibular nucleus; MVeMC, medial vestibular nucleus magnocellular part; Gi, gigantocellular nucleus; MdV, ventral medial reticular nucleus; PMn, paramedian reticular nucleus; GiA, anterior gigantocellular nucleus; GiV, ventral gigantocellular nucleus; LPGi, lateral paragigantocellular nucleus; LRt, lateral reticular nucleus; RMg, raphe magnus; ROb, raphe obscuruis; RPa, raphe pallidus (Paxinos & Franklin, 2004).
